# Alpha-Frag: a deep neural network for fragment presence prediction improves peptide identification

**DOI:** 10.1101/2021.04.07.438629

**Authors:** Jian Song, Fangfei Zhang, Changbin Yu

## Abstract

Identification of peptides in mass spectrometry-based proteomics typically relies on spectra matches. As MS/MS spectra record presence and intensity of fragment ions, the match should take both fragment presence similarity and intensity similarity into consideration. Fragment presence similarity can be calculated with the help of fragment presence prediction such as theoretical enumeration of all possible fragment ions or selecting non-zero intensity ions from the result of fragment intensity prediction, but neither of these two methods is accurate enough. In this work, we developed a deep neural network based model, Alpha-Frag, to predict precisely the fragment ions that should be present for a given peptide. Alpha-Frag modelled fragment presence prediction as a multi-label classification task and trained with ProteomeTools dataset. In terms of intersection over union (IoU), Alpha-Frag achieved an average of >0.7 and outperformed the benchmarks across the validation datasets. Furthermore, fragment presence similarity was calculated based on presence prediction and incorporated into the peptide statistical validation tools as an additional score to improve peptide identifications. Our preliminary experiments show that this score led to a maximum increase of 26.8% (FDR 0.1%) and 21.6% (FDR 1%) for the DDA and the DIA identification, respectively.

**Significance Statement:** A better prediction of fragmentation for peptides in mass spectrometry (MS) is beneficial to the peptide identification. As the MS/MS spectra record two-dimensional information of fragment ions derived from precursors, mass-to-charge ratio (m/z) and their corresponding intensities, besides the fragment intensity prediction, it is necessary to study the presence prediction. Although the presence prediction can be realized by enumerating all the possible fragmentation patterns of a peptide with equal probability or by selecting non-zero intensity fragment ions from the result of fragment intensity prediction, neither of these two methods is accurate enough. In this study, deep learning is leveraged to precisely predict the fragment ions of a given peptide. Based on the fragment presence prediction, fragment presence similarity between experimental spectra and predicted spectra can be calculated which is proved to promote the peptide detections both for DDA and for DIA data.

## 1 Introduction

Tandem mass spectrometry (MS/MS)-based technologies have been widely used in bottom-up proteomics for peptide identification and quantification [1]. The MS/MS spectra record two-dimensional information of fragment ions derived from precursors, mass-to-charge ratio (*m/z*) and their corresponding intensities. Data-dependent acquisition (DDA) and data-independent acquisition (DIA) are the two most prevalent MS/MS strategies. Identification of peptides in DDA mainly relies on the match between the experimental spectrum and the theoretical spectrum. The ratio of the number of observed ions to the number of all theoretically possible fragment ions is an import evidence to the peptide-to-spectrum match (PSM) [2,3]. For DIA, identification of peptides is heavily based on fragment intensity match between the extracted ion chromatograms (XIC) and the reference spectral library which contains a certain number of most intensive fragment ions produced by DDA [4,5]. As MS/MS spectra record the presence and intensity of fragment ions simultaneously, the match should take both fragment presence similarity and intensity similarity into consideration both for DDA and for DIA identification (Similar to the term fragment intensity, which describes the intensities of fragment ions, the term fragment presence means the collection of fragment ions regardless of their intensities. Further, fragment presence similarity represents the ratio of the number of the matched-predicted ions to the number of the predicted fragment ions). When more realistic set of fragment ions rather than theoretical possible fragment ions is provided to calculate fragment presence similarity, the evidences of DDA and DIA identification will be more accurate and more comprehensive, respectively.

The presence and absence of fragment ions observed in an MS/MS spectrum depend on factors including peptide primary sequence, charge state of the peptide, and how the collision energy is introduced [6]. To achieve fragment presence prediction, enumerating all the possible fragmentation patterns of a peptide with equal probability is possible theoretically but leads to produce a lot of redundant fragment ions. Alternatively, selecting non-zero intensity fragment ions from the result of fragment intensity prediction is another implementation of fragment presence prediction. Currently, with the advances in deep learning based modeling [7], fragment intensity predictors modeled by neutral network, such as pDeep [8], DeepMass [9] and Prosit [10], are making increasingly accurate fragment intensity prediction. Among these, pDeep and DeepMass followed the framework of bidirectional LSTM [11] combined with fully connected layers to predict intensities of fragment ion. Innovatively, Prosit took advantage of the recent advances in neural machine translation [12,13] with an encoder to code the amino acids sequence and a decoder to output the fragment ion intensity step by step and had become the state-of-art of intensity predictor. For fragment presence prediction based on intensity predictors, the above-mentioned intensity prediction models aim to spectrum similarities [14] which dominated by high-intensity ions, potentially limiting existence prediction of low-intensity ions. Some low-intensity fragment ions that non-existing but predicting present or present but predicting missing may lead to inaccurate calculation of presence similarity and reduce subsequent statistical power for peptide identification.

The purpose of this study is to develop a deep neural network-based model, Alpha-Frag, for fragment presence prediction. Our study design, including model training, evaluation and applications, is shown in Fig. 1. Peptide sequence and charge are taken as the input for Alpha-Frag to predict the presence of fragments ions. Unlike intensity prediction, which is a regression problem in most cases, Alpha-Frag models fragment presence prediction as a multi-label classification problem. Taking advantage of the ProteomeTools [15] dataset for model training, Alpha-Frag can output the most likely set of fragment ions. The result of prediction was evaluated by intersection over union (IoU) between experimental and predicted fragment ions. To illustrate the possible applications of Alpha-Frag, we calculated the ratio of the number of matched-predicted ions versus the number of predicted ions by Alpha-Frag as an additional score to improve DDA and DIA identification. The preliminary results showed that Alpha-Frag is able to predict the presence of fragment ions accurately and is beneficial to both DDA and DIA identifications.

**Fig. 1.**
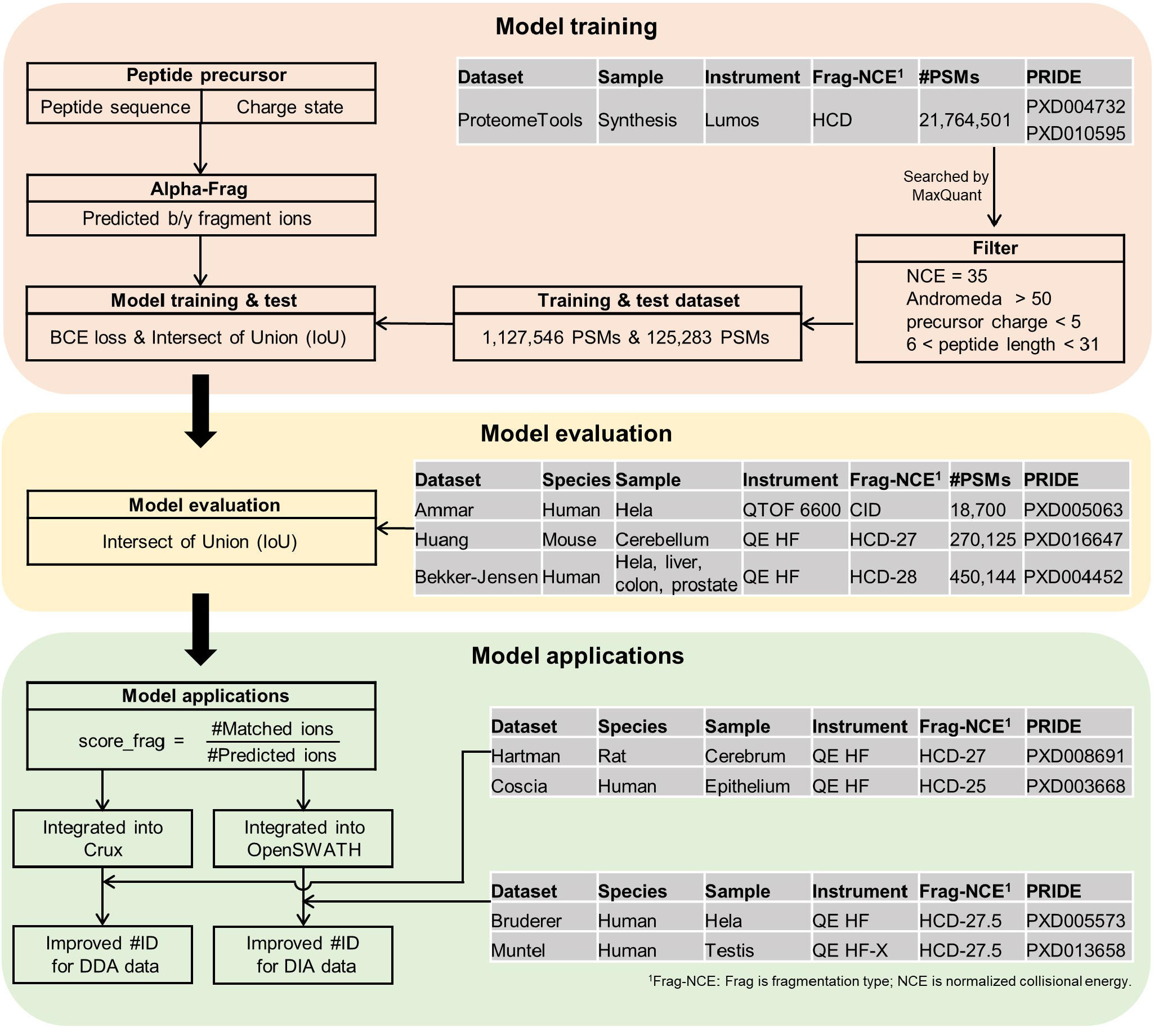
Study design. From top to bottom, the three panels indicate model training, model evaluation and model applications, respectively. The tables in each panel are the corresponding datasets.

## 2 Materials and methods

### 2.1 Data preparation

In this paper, multiple datasets were used to model training & test, evaluation and applications (shown in Fig. 1). Like Prosit, training and test datasets (the table in top panel of Fig. 1) were sourced from ProteomeTools (PXD004732 and PXD010595 in PRIDE) of which the raw data were searched by MaxQuant [16] (v.1.5.3.30) at 1% peptide false discovery rate (FDR). To select a high confident dataset for training and test, only PSMs met these criteria were considered: Andromeda score is >50, charge is <5, peptide length is restricted to 7-30 amino acids. Alpha-Frag used MS/MS data with normalized collision energy (NCE) 35 suggested by [17]. Meanwhile, y/b fragment ions with charges up to 2 were extracted and annotated from the corresponding MS/MS for each identified peptide by MaxQuant, and these y/b ions were used as the ground truth of present fragment ions for training and test. In total, 452,558 precursors (1,252,829 PSMs) were kept and split into a training set and a test set randomly by a ratio at 4:1.

To ensure the generality of model evaluation, we relied on three diverse DDA datasets. Details were summarized in the table in the middle panel of Fig. 1. Among these, Ammar’s dataset [18] was acquired by TripleTOF 6600 with CID fragmentation. After searched by MaxQuant (v.1.5.3.12) at 1% FDR and filtration (Andromeda score is >50, precursor charge is <5, peptide length is restricted to 7-30 amino acids), Ammar’s dataset has 18,700 PSMs. Huang’s dataset [19] was obtained from Q Exactive HF (Thermo Scientific) with HCD fragmentation (NCE=27) and has 270,125 PSMs after the same filtration (MaxQuant v.1.6.0.1). Bekker-Jensen’s dataset [20] searched by MaxQuant (v.1.5.3.6) was conducted in Q Exactive HF with HCD fragmentation (NCE=28) and has 450,144 PSMs after merged search results. Each of the PSMs for these three datasets had a corresponding set of present fragment ions determined by MaxQuant. For training & test and evaluation datasets, the search result of MaxQuant were available on their corresponding PRIDE.

For the model application in DDA identification, Hartman’s dataset [21] and Coscia’s dataset [22] both without fractionation were adopted to retain sample complexity (the table in button panel of Fig. 1). These two datasets were acquired by Q Exactive HF with NCE 27 and 25, respectively. In order to facilitate the integration of fragment presence prediction to DDA identification, search software Crux [3] and Percolator [23] were selected. The acquired RAW data were analyzed with Crux (v.3.2) and Percolator (v.3.02.0) against the rat UniProt fasta database (state 08.24.2020, 29,940 entries) and the human SwissProt fasta database (state 09.03.2020, 20,379 entries) for Hartman’s and Coscia’s dataset, respectively. The options of Crux included: ‘--min-length 7 --max-length 30 --mods-spec C+57.02146,1M+15.994915 –max-mods 3 –missed-cleavages 1 --max-precursor-charge 4 --top-match 1 --concat T --compute-sp T’. Percolator used the default parameters.

For the application of Alpha-frag in DIA identification, Bruderer’s dataset [24] and Muntel’s dataset [25] were adopted to represent the cell line (HeLa) and the complex clinical tissue sample (testis of human), respectively. Bruderer’s dataset was collected on a Q Exactive HF with 37 isolation windows and Muntel’s dataset was acquired on a Q Exactive HF-X (Thermo Scientific) with 45 isolation windows. Both Bruderer’s and Muntel’s dataset had an NCE of 27.5. Software OpenSWATH [5] and PyProphet [26] were selected to identify the DIA files as they are convenient to add custom function. Meanwhile, the PHL public spectral library [27] which contains target and decoy peptides and the CiRT peptides [28] were used to aid the identification. The following options were employed for OpenSWATH (v.2.4.0) : ‘-batchSize 1000 -readOptions cacheWorkingInMemory-mz_extraction_window 20 -ppm -mz_correction_function quadratic_regression_delta_ppm - min_rsq 0.8’. PyProphet (v.0.24.1) used the default parameters.

For all dataset in this study, only peptides with fixed modification of carbamidomethylation on cysteine and variable modification of oxidation on methionine were considered.

### 2.2 Model

For fragment presence prediction based on peptide sequence, the predictor should be able to capture the effects of all amino acids on the fragment result and be able to handle sequential input with a variable length. Similar to Prosit, Alpha-Frag also uses bi-directional gated recurrent memory units (Bi-GRU) [29] to address these requirements. Fig. 2a is the architecture overview of Alpha-Frag. A detailed description of the model is presented below.

**Fig. 2.**
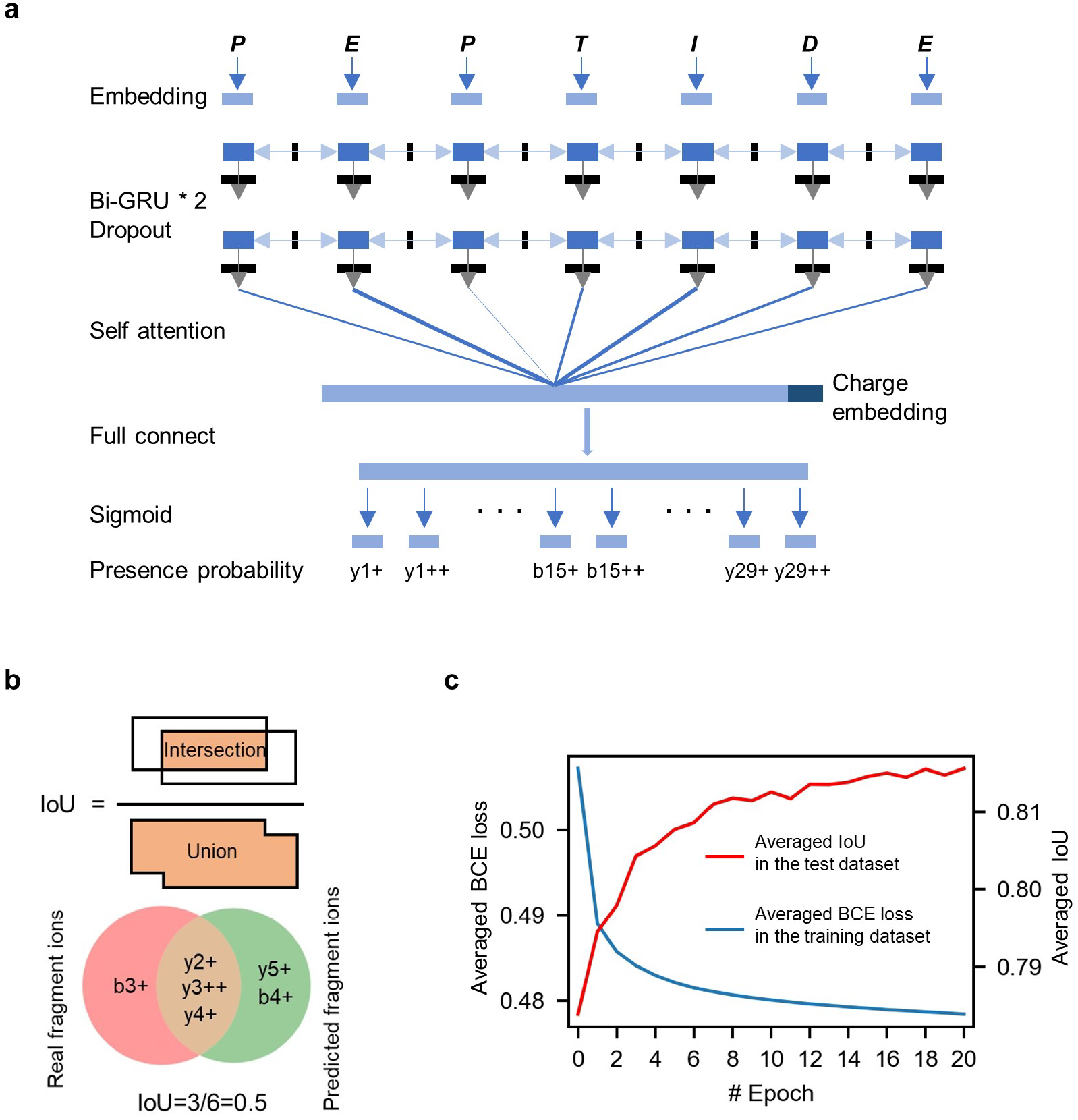
The model of Alpha-Frag and its training and test. **a)** Bi-GRU based Alpha-Frag model. **b)** Illustration of intersection over union (IoU) which is used to compare the consistence between two sets. **c**) The averaged loss curve in training dataset and the averaged IoU curve in test dataset.

#### Inputs

Including peptide sequence and precursor charge. Each amino acid of peptide and precursor charge will be embedded to a vector of dimension 32. Alpha-Frag supports precursor charge with no more than 4, as well as 20 natural amino acids and variable modification of oxidation on methionine. Each cysteine is treated as fixed modification of carbamidomethylation on itself.

#### Model

Alpha-Frag connects embedding layer to two layers of Bi-GRU. Dropout [30] is set to 0.5 between the two Bi-GRU layers. A self-attention layer [31] is used to merge the Bi-GRU outputs to a vector of dimension 64. Concatenating this vector with charge embedding vector, Alpha-Frag represents the precursor as a vector of dimension 96 in latent space. Afterwards, the 96-dimensional vector is converted to dimension of 116 (Alpha-Frag considers peptide length maximum to 30 and only y/b fragment ions with no more than 2 charges. The maximum number of fragment ions is: 2 * 2 * 29 = 116) by a fully connected layer and a sigmoid layer.

#### Outputs

The output 116-dimensional vector is ordered as follows: y1+, y1++, b1+, b1++, y2+ and so on. Each value represents the occurrence probability of the corresponding fragment ion. Alpha-Frag adopts 0.5 as the existence threshold. Only fragment ion with a predicted probability greater than 0.5 will be considered as presence. Fragment ions at impossible dimensions (such as y17 for an 8-mer peptide) are set to zero directly.

Of note, we experimented with different dropout and dimension of each layer. However, this did not result in any significant gains on the performance. Therefore, we used the common parameter values described above. The source code of Alpha-Frag is available at https://github.com/YuAirLab/Alpha-Frag.

### 2.3 Training and test

PyTorch (v.1.1.0, https://pytorch.org) was used to implement and train Alpha-Frag. We used the Adam optimizer with an initial learning rate of 0.001 and 1024 samples per batch. The loss function in training was binary cross entropy loss (BCE loss):

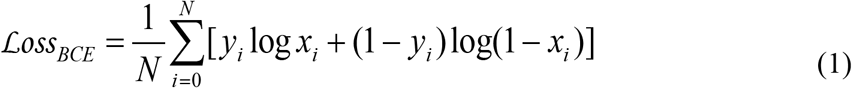

where *x*_*i*_ means the true label of the corresponding fragment ion either 0 or 1, *y*_*i*_ is the predicted probability, and *N* is the length of the output vector. It should be noted that the fragment ions out-of-range or out-of-charge were masked and not involved in the calculation of loss. For the test dataset, we calculated the intersection over union (IoU, Fig. 2b) to evaluate the accuracy of prediction. Given a set of real present fragment ions **𝒜** and a set of predicted present fragment ions **ℬ**, the IoU is defined as follows:

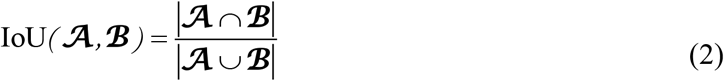

For example, the real fragment ions is ‘y2+, y3+, y4+, y5++, b2+, b3+’, the predicted fragment ions is ‘y1+, y2+, y4+, y5+, b3+’, then the intersection between them is ‘y2+, y4+, b3+’ and the union is ‘y1+, y2+, y3+, y4+, y5+, y5++, b2+, b3+’. Hence, the IoU value is equal to 3/8. The larger the IoU, the more consistent the predicted fragment ions with the real fragment ions are.

## 3 Results

### 3.1 Bi-GRU based deep learning model

We trained Alpha-Frag on one Nvidia 1060 GPU. For each epoch, we calculated the averaged BCE loss and the averaged IoU in the training and test dataset, respectively. As the epoch increased, the averaged BCE loss decreased and the averaged IoU increased shown in Fig. 2c. After 20 epochs, the model converged and the training was terminated to balance the learning and the potential overfitting. At last, the averaged IoU reached 0.815, indicating that Alpha-Frag predict the present fragment ions well for the test dataset.

### 3.2 Performance Evaluation of fragment presence prediction

In order to illustrate fragment presence prediction based on intensity prediction is not accurate enough, we plotted the difference of a representative precursor EPPLLLGVLHPNTK(3+) between its experimental spectrum, the predicted spectrum by Prosit (unless otherwise specified, Prosit works with the optimal NCE 35 in this study) and its predicted fragment ions by Alpha-Frag in Fig. 3a. Although the Pearson correlation coefficient (PCC) between the experimental spectrum and the predicted spectrum by Prosit reached as high as 0.90, the predicted spectrum contained 3.06 (46 vs. 15) times as many fragment ions as the number of matched y/b ions in the experimental spectrum. In other words, ∼67.3% of predicted fragment ions by Prosit are not present. As a contrast, Alpha-Frag predicted 14 fragment ions and 13 of which are matched to experimental spectrum and one of which is mismatch. Therefore, the IoU between the real present fragment ions and the predicted present fragment ions by Alpha-Frag reached 0.81, illustrating the strong agreement between prediction and observation for fragment presence in this case.

**Fig. 3.**
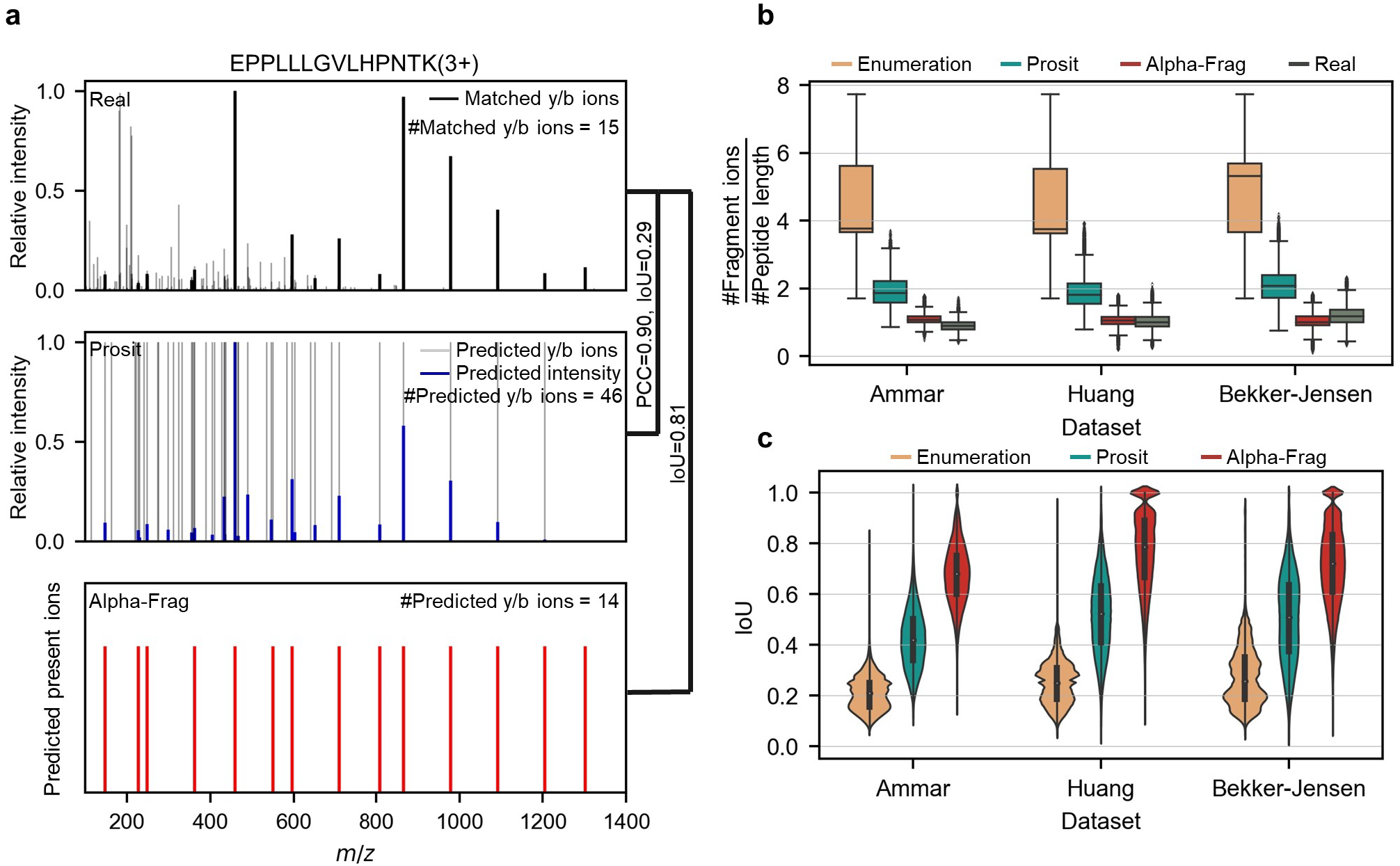
Performance evaluation. **a)** Comparison between experimental spectrum, predicted spectrum by Prosit (NCE=35), predicted fragment ions by Alpha-Frag of precursor EPPLLLGVLHPNTK(3+). The matched y/b ions in experimental spectrum contains 15 y/b ions marked in jet black lines (scan 53773, 01974C_BA1-TUM_missing_first_1_01_01-DDA-1h-R4.raw, PXD010595. The matched y/b ions are determined by MaxQuant at 1% FDR). The predicted spectrum by Prosit contains 46 y/b ions with non-zero intensities marked in light black. The blue lines mean the match between the predicted spectrum and the experimental spectrum. We observed that 67.3% of predicted fragment ions are not present in the experimental spectrum while the Pearson correlation coefficient (PCC) between them reaches 0.90. In the other hand, the IoU between experimental spectrum and predicted fragment ions by Alpha-Frag reaches 0.81. **b)** Box plot to compare the predicted number of fragment ions and the real number of fragment ions both normalized by peptide length. The box indicates the interquartile range (IQR), its whiskers 1.5× IQR values, and the black line the median. **c)** Violin plot of IoU in three evaluation datasets.

To evaluate the performance of fragment presence prediction of Alpha-Frag comprehensively, we benchmarked it against theoretical enumeration model (hereinafter referred to as Enumeration model) and Prosit model using three public DDA datasets: Ammar’s, Huang’s and Bekker-Jensen’s dataset. For Enumeration model, the predicted fragment ions include all possible y/b ions with less length and no more charges than precursors. First, we took a rough look at the difference between the predicted number of fragment ions and the real number of fragment ions both normalized by peptide length in Fig. 3b. In all three datasets, the result of Enumeration model and Prosit model overestimated the number of fragment ions and Alpha-Frag was the most closest model compared to the real number. This comparison showed that Alpha-Frag is able to predict precisely the number of fragment ions.

Besides the above comparison, IoU can be calculated to check the consistence between the real fragment ions and the predicted fragment ions. Fig. 3c plotted the IoU distribution by violin plot. It is apparent that Alpha-Frag outperformed markedly Enumeration model and Prosit model across the datasets. In terms of median IoU, Alpha-Frag achieved an average of ∼207% and ∼52% improvements than Enumeration model and Prosit model, respectively, indicating that Alpha-Frag works very well for fragment presence prediction.

Last but not least, in our device of CPU Intel i7-7700K 4 cores, Win10, 64 bit, 64 GB memory and GPU Nvidia GTX 1060, Enumerator model, Prosit and Alpha-Frag spent ∼0s, ∼32.5s and ∼5.2s to predict fragment presence of 450,144 precursors from Berken-Jensen’s dataset, respectively.

### 3.3 Presence prediction improves peptide identification for both DDA and DIA

To test if the presence prediction is beneficial to peptide identification, we constructed an additional score, termed *score_frag*, and integrated it into the corresponding pipeline (default 20 ppm tolerance for matching):

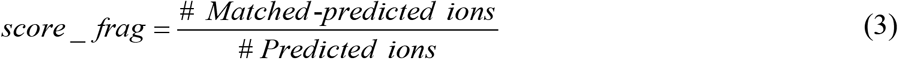

For DDA identification (Fig. 4a), Crux and Percolator were employed to scored (such as Sp and XCorr) each candidate PSM. Then, fragment presence predictor was used to generate the predicted fragment ions based on the peptide sequence and precursor charge. Subsequently, the *score_frag* was calculated based on the real spectrum and the predicted fragment ions and was fed into Percolator with the other scores (as shown by blue lines in Fig. 4a). At last, Percolator merged all scores and calculated the FDR. We carried out this DDA *score_frag* strategy with Enumeration model, Prosit model and Alpha-Frag to two DDA datasets: Hartmann’s dataset and Coscia’s dataset. Raw identification without *score_frag* was used as a control. The number of identified PSMs against FDR was shown in Fig. 4b as well as the bar plot of scores’ weight (the Alpha-Frag case). Overall, we found that including *score_frag* increases the number of identified PSMs compared to raw identification. In terms of the identification number, Alpha-Frag outperformed Enumeration model and Prosit model at the stringent FDR range (< 0.5%). While along FDR to 5%, no significant differences were found between Alpha-Frag and Prosit (data not shown). At 0.1% FDR, raw identification reached only an average of 80.3% and 79.6% compared to Alpha-Frag for Hartmann’s and Coscia’s dataset, respectively. Meanwhile, the weight of *score_frag* was ranked number one across the datasets, illustrating that *score_frag* based on fragment presence prediction is an important evidence for DDA identification.

**Fig. 4.**
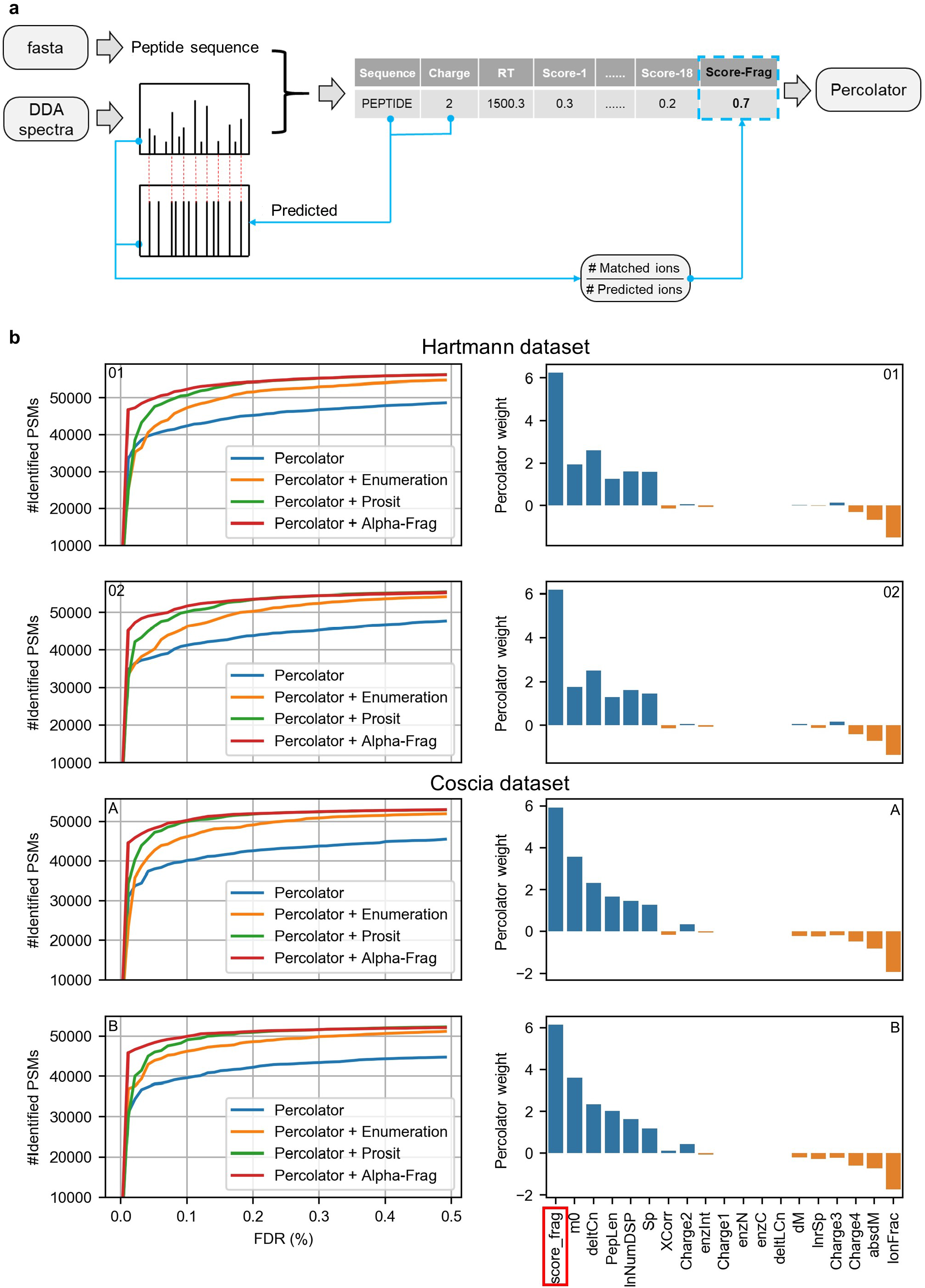
Application of fragment presence prediction to DDA identification. **a)** Workflow. The non-colored lines represent the workflow of DDA identification by Crux combined with Percolator. The blue lines represent the process of integrating the *score_frag* into Crux-Percolator. **b)** Comparison of identification performance and weight of scores. In Hartmann’s dataset, ‘01’ refers to 20160809_EXQ00_DaHo_SA_Eddie_15mioPCN_fullproteome_2016_06_07_GFP_01.raw, ‘02’ refers to 20160809_EXQ00_DaHo_SA_Eddie_15mioPCN_fullproteome_2016_06_07_GFP_02.raw. In Coscia’s dataset, ‘A’ refers to 20130312_EXQ4_FaCo_SA_CL_DOV13_A.raw, ‘B’ refers to 20130312_EXQ4_FaCo_SA_CL_DOV13_B.raw. Right side is the bar plot of Percolator weights for the ‘Percolator + Alpha-Frag’ case.

Application of *score_frag* to DIA identification is different from that to DDA identification. As shown in Fig. 5a, after OpenSWATH determined and scored the peak groups based on the spectra library, fragment presence predictor was performed to predict the fragment presence ions based on the peptide sequence and precursor charge. Then, depending on the apex retention time of the peak group, the single MS/MS spectrum was picked out whose retention time was closest to the apex. Afterward, the *score*_*frag* was calculated based to the picked spectrum and the predicted fragment ions and was fed into PyProphet with the other scores (blue lines in Fig. 5a). At last, PyProphet merged the scores and reported the identification result. Similarly, we implemented this DIA *score_frag* strategy with Enumeration model, Prosit model and Alpha-Frag model to two DIA datasets: Bruderer’s and Muntel’s dataset. As a control, we also analyzed data without *score_frag*.

**Fig. 5.**
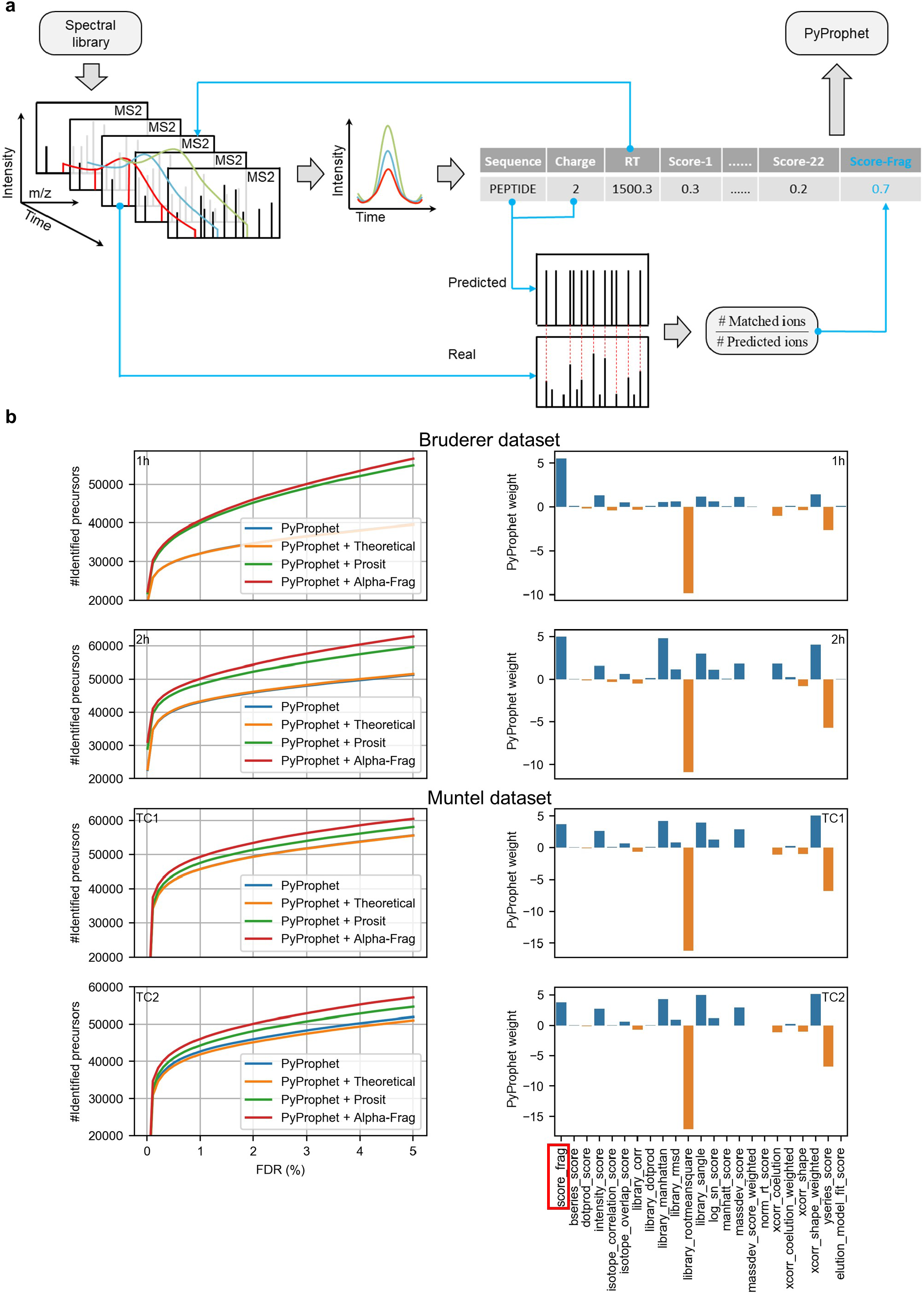
Application of fragment presence prediction to DIA identification. **a)** Workflow. The non-colored lines represent the workflow of DIA identification by OpenSWATH combined with PyProphet. The blue lines represent the process of integrating the *score_frag* into OpenSWATH-PyProphet. **b)** Comparison of identification performance and weight of scores. In Bruderer’s dataset, ‘1h’ refers to Fig2 HeLa-1h_MHRM_R01_T0.raw, ‘2h’ refers to Fig2 HeLa-2h_MHRM_R01_T0.raw. In Muntel’s dataset, ‘TC1’ refers to G_D190415_S553-TestisCancerSet-2h-2ug-TC1_MHRM_R01_T0.raw, ‘TC2’ refers to G_D190415_S553-TestisCancerSet-2h-2ug-TC2_MHRM_R01_T0.raw. Right side is the bar plot of PyProphet weights for the ‘PyProphet + Alpha-Frag’ case.

The number of identified precursors against the FDR was plotted in Fig. 5b as well as the scores’ weight (the Alpha-Frag case). In terms of the number of identified precursors, we found that Enumeration model did not always lead to improvement. This may be due to the fact that the MS/MS in DIA is multiplexed. Besides, in all datasets, Alpha-Frag outperformed Prosit model, both of which increased the identification. At 1% FDR, Alpha-Frag brought about an average of 21.6% and 7.9% improvement over the raw identification for Bruderer’s and Muntel’s dataset, respectively.

Dissimilar in DDA, the weight of *score_frag* in DIA was not ranked number one but had a same order of magnitude compared to the most important score. These results demonstrated the feasibility of the additional *score_frag* to DIA identification and the accuracy of fragment presence prediction by Alpha-Frag.

## 4 Concluding remarks

In this study, we proposed Alpha-Frag, a deep neural network used to predict fragment presence ions and that surpass substantially the other benchmarks. In the validation datasets, Alpha-Frag achieved an average of ∼207% and ∼52% improvements in terms of median IoU than Enumeration model and Prosit model, respectively. This highlights that the model is an effective complement to fragment intensity prediction. In the accompanying applications of Alpha-Frag, fragment presence similarity was performed to improve the identification and achieved a maximum increase of 26.8% (FDR 0.1%) and 21.6% (FDR 1%) for the DDA dataset and the DIA dataset, respectively. The weight of fragment presence similarity also indicates that it is an import evidence for peptide identification. So far, only tryptic peptides and peptides with no modifications except for methionine oxidation were considered. Next as more data for non-tryptic and modified peptides and chemical labels become available, we will update our model to improve predictions for more peptides. Further, we plan to divide the spectrum prediction into two phases: fragment presence prediction followed by corresponding intensity prediction to approximate the fragmentation better. We believe this will make the predicted spectrum more accurate and peptide identification more confident both for DDA and for DIA.

## Acknowledgements

This study was funded by National Natural Science Foundation of China (grant number 82003766, 31970636), Academic Promotion Project of Shandong First Medical University.

## References

1. Aebersold, R., & Mann, M. (2016). Mass-spectrometric exploration of proteome structure and function. Nature, 537(7620), 347–355.

2. Eng, J. K., McCormack, A. L., & Yates, J. R. (1994). An approach to correlate tandem mass spectral data of peptides with amino acid sequences in a protein database. Journal of the american society for mass spectrometry, 5(11), 976–989.

3. McIlwain, S., Tamura, K., Kertesz-Farkas, A., Grant, C. E., Diament, B., Frewen, B., … & Noble, W. S. (2014). Crux: rapid open source protein tandem mass spectrometry analysis. Journal of proteome research, 13(10), 4488–4491.

4. Gillet, L. C., Navarro, P., Tate, S., Röst, H., Selevsek, N., Reiter, L., … & Aebersold, R. (2012). Targeted data extraction of the MS/MS spectra generated by data-independent acquisition: a new concept for consistent and accurate proteome analysis. Molecular & Cellular Proteomics, 11(6).

5. Röst, H. L., Rosenberger, G., Navarro, P., Gillet, L., Miladinović, S. M., Schubert, O. T., … & Aebersold, R. (2014). OpenSWATH enables automated, targeted analysis of data-independent acquisition MS data. Nature biotechnology, 32(3), 219–223.

6. Perkins, D. N., Pappin, D. J., Creasy, D. M., & Cottrell, J. S. (1999). Probability - based protein identification by searching sequence databases using mass spectrometry data. ELECTROPHORESIS: An International Journal, 20(18), 3551–3567.

7. LeCun, Y., Bengio, Y., & Hinton, G. (2015). Deep learning. nature, 521(7553), 436–444.

8. Zhou, X. X., Zeng, W. F., Chi, H., Luo, C., Liu, C., Zhan, J., … & Zhang, Z. (2017). pDeep: predicting MS/MS spectra of peptides with deep learning. Analytical chemistry, 89(23), 12690–12697.

9. Tiwary, S., Levy, R., Gutenbrunner, P., Soto, F. S., Palaniappan, K. K., Deming, L., … & Cox, J. (2019). High-quality MS/MS spectrum prediction for data-dependent and data-independent acquisition data analysis. Nature methods, 16(6), 519–525.

10. Gessulat, S., Schmidt, T., Zolg, D. P., Samaras, P., Schnatbaum, K., Zerweck, J., … & Wilhelm, M. (2019). Prosit: proteome-wide prediction of peptide tandem mass spectra by deep learning. Nature methods, 16(6), 509–518.

11. Hochreiter, S., & Schmidhuber, J. (1997). Long short-term memory. Neural computation, 9(8), 1735–1780.

12. Kingma, D. P., & Ba, J. (2014). Adam: A method for stochastic optimization. arXiv preprint 1412.6980.

13. Xu, K., Ba, J., Kiros, R., Cho, K., Courville, A., Salakhudinov, R., … & Bengio, Y. (2015, June). Show, attend and tell: Neural image caption generation with visual attention. In International conference on machine learning (pp. 2048–2057). PMLR.

14. Toprak, U. H., Gillet, L. C., Maiolica, A., Navarro, P., Leitner, A., & Aebersold, R. (2014). Conserved peptide fragmentation as a benchmarking tool for mass spectrometers and a discriminating feature for targeted proteomics. Molecular & Cellular Proteomics, 13(8), 2056–2071.

15. Zolg, D. P., Wilhelm, M., Schnatbaum, K., Zerweck, J., Knaute, T., Delanghe, B., … & Kuster, B. (2017). Building ProteomeTools based on a complete synthetic human proteome. Nature methods, 14(3), 259–262.

16. Cox, J., & Mann, M. (2008). MaxQuant enables high peptide identification rates, individualized ppb-range mass accuracies and proteome-wide protein quantification. Nature biotechnology, 26(12), 1367–1372.

17. Zolg, D. P., Wilhelm, M., Yu, P., Knaute, T., Zerweck, J., Wenschuh, H., … & Kuster, B. (2017). PROCAL: a set of 40 peptide standards for retention time indexing, column performance monitoring, and collision energy calibration. Proteomics, 17(21), 1700263.

18. Ammar, C., Berchtold, E., Csaba, G., Schmidt, A., Imhof, A., & Zimmer, R. (2019). Multi-reference spectral library yields almost complete coverage of heterogeneous LC-MS/MS data sets. Journal of proteome research, 18(4), 1553–1566.

19. Huang, T., Bruderer, R., Muntel, J., Xuan, Y., Vitek, O., & Reiter, L. (2020). Combining precursor and fragment information for improved detection of differential abundance in data independent acquisition. Molecular & Cellular Proteomics, 19(2), 421–430.

20. Bekker-Jensen, D. B., Kelstrup, C. D., Batth, T. S., Larsen, S. C., Haldrup, C., Bramsen, J. B., … & Olsen, J. V. (2017). An optimized shotgun strategy for the rapid generation of comprehensive human proteomes. Cell systems, 4(6), 587–599.

21. Hartmann, H., Hornburg, D., Czuppa, M., Bader, J., Michaelsen, M., Farny, D., … & Edbauer, D. (2018). Proteomics and C9orf72 neuropathology identify ribosomes as poly-GR/PR interactors driving toxicity. Life science alliance, 1(2).

22. Coscia, F., Watters, K. M., Curtis, M., Eckert, M. A., Chiang, C. Y., Tyanova, S., … & Mann, M. (2016). Integrative proteomic profiling of ovarian cancer cell lines reveals precursor cell associated proteins and functional status. Nature communications, 7(1), 1–14.

23. Käll, L., Canterbury, J. D., Weston, J., Noble, W. S., & MacCoss, M. J. (2007). Semi-supervised learning for peptide identification from shotgun proteomics datasets. Nature methods, 4(11), 923–925.

24. Bruderer, R., Bernhardt, O. M., Gandhi, T., Xuan, Y., Sondermann, J., Schmidt, M., … & Reiter, L. (2017). Optimization of experimental parameters in data-independent mass spectrometry significantly increases depth and reproducibility of results. Molecular & Cellular Proteomics, 16(12), 2296–2309.

25. Muntel, J., Gandhi, T., Verbeke, L., Bernhardt, O. M., Treiber, T., Bruderer, R., & Reiter, L. (2019). Surpassing 10000 identified and quantified proteins in a single run by optimizing current LC-MS instrumentation and data analysis strategy. Molecular omics, 15(5), 348–360.

26. Teleman, J., Röst, H. L., Rosenberger, G., Schmitt, U., Malmström, L., Malmström, J., & Levander, F. (2015). DIANA—algorithmic improvements for analysis of data-independent acquisition MS data. Bioinformatics, 31(4), 555–562.

27. Rosenberger, G., Koh, C. C., Guo, T., Röst, H. L., Kouvonen, P., Collins, B. C., … & Aebersold, R. (2014). A repository of assays to quantify 10,000 human proteins by SWATH-MS. Scientific data, 1(1), 1–15.

28. Parker, S. J., Rost, H., Rosenberger, G., Collins, B. C., Malmström, L., Amodei, D., … & Aebersold, R. (2015). Identification of a Set of Conserved Eukaryotic Internal Retention Time Standards for Data-independent Acquisition Mass Spectrometry*[S]. Molecular & Cellular Proteomics, 14(10), 2800–2813.

29. Chung, J., Gulcehre, C., Cho, K., & Bengio, Y. (2014). Empirical evaluation of gated recurrent neural networks on sequence modeling. arXiv preprint 1412.3555.

30. Srivastava, N., Hinton, G., Krizhevsky, A., Sutskever, I., & Salakhutdinov, R. (2014). Dropout: a simple way to prevent neural networks from overfitting. The journal of machine learning research, 15(1), 1929–1958.

31. Lin, Z., Feng, M., Santos, C. N. D., Yu, M., Xiang, B., Zhou, B., & Bengio, Y. (2017). A structured self-attentive sentence embedding. arXiv preprint 1703.03130.

